# Reward contingency modulates olfactory bulb output via pathway-dependent peri-somatic inhibition

**DOI:** 10.1101/2023.08.17.553686

**Authors:** Sander Lindeman, Xiaochen Fu, Janine K Reinert, Izumi Fukunaga

## Abstract

Associating values to environmental cues is a critical aspect of learning from experiences, allowing animals to predict and maximise future rewards. Value-related signals in the brain were once considered a property of higher sensory regions, but its wide distribution across many brain regions is increasingly recognised. Here, we investigate how reward-related signals begin to be incorporated, mechanistically, at the earliest stage of olfactory processing, namely, in the olfactory bulb. In head-fixed mice performing Go/No-Go discrimination of closely related olfactory mixtures, rewarded odours evoke widespread inhibition in one class of output neurons, that is, in mitral cells but not tufted cells. The temporal characteristics of this reward-related inhibition suggest it is odour-driven, but it is also context-dependent since it is absent during pseudo-conditioning and pharmacological silencing of the piriform cortex. Further, the reward-related modulation is present in the somata but not in the apical dendritic tuft of mitral cells, suggesting an involvement of circuit component located deep in the olfactory bulb. Depth-resolved imaging from granule cell dendritic gemmules suggests that granule cells that target mitral cells receive a reward-related extrinsic drive. Our results support the notion that value-related modulation appears at the early stages of sensory processing and provide constraints on long-range and local circuit mechanisms.

## Introduction

Sensory systems of the brain play a crucial role in guiding animals’ choices. One of their uses is in the reward-driven decision-making, where the system is thought to adjust the representations of sensory cues depending on the past reward encounters, to influence their future behavioural choices and learning. Decades of studies across brain areas have demonstrated that reward expectations are, in turn, potent modulators of sensory activity. For example, stimulus evoked responses in many sensory regions of the brain scale with the quantity of expected reward ^1–5^. This modulation is often interpreted as representing subjective value ^6,7^. Such a system where sensory processing is fined-tuned flexibly may be crucial for maximising returns in a dynamic and uncertain world ^7^.

Decision and value-related modulations of sensory responses are featured prominently in higher sensory areas ^1,8^. However, recent studies indicate that even early stages of sensory processing, especially in rodents, participate in value-like representations, showing ample modulations associated with decision-making ^9–11^. The olfactory system is an extreme case in this regard, where apparent reward-related modulation is readily observed as peripherally as in the olfactory bulb ^12,13^, the primary olfactory region situated just one synapse away from the site of sensory transduction. This peripheral location, along with the saliency of the olfactory cues for rodents, makes the olfactory bulb an attractive structure to study the mechanisms that generate value-like signals in the brain ^12^.

The nature of this apparent reward-related modulation in the olfactory bulb remains unresolved. For example, one study observed that evoked responses to rewarded vs. unrewarded odours in the principal neurons of the olfactory bulb diverge only transiently as rats learn to discriminate between the cues ^12^. This may reflect the level of animal’s engagement ^14^, where the learning-related modulation corresponds mainly to the changes in the inputs from the sensory periphery arising from sniff pattern changes. Rodents indeed adjust the odour sampling patterns exquisitely according to the behavioural contexts ^15,16^. However, given that the olfactory bulb is a major target of feedback and neuromodulatory projections from many brain regions, value-related information could affect how the olfactory bulb represents odours. For example, electrical and optogenetic stimulations and pharmacological manipulations of neuromodulatory and feedback inputs to the olfactory bulb change the gain of odour responses in the principal neurons of this region ^17–22^. Therefore, whether the reward-related signals in the primary olfactory area simply reflect changes in the sensory input or internally generated contextual signals need to be clarified.

Here, we show that the olfactory bulb exhibits robust and consistent reward-related signals during a trace olfactory conditioning paradigm, where mice discriminate between closely related olfactory mixtures. This phenomenon is characterised by widespread inhibitory responses following the rewarded odour presentation, in mitral cells but not tufted cells. This divergence is not explained by the odour identity or sampling strategy and reflects the congruence of sensory drive and contextual signals. By imaging from specific subcellular compartments of mitral cells, we demonstrate that the divergent responses first become evident peri-somatically. Depth-resolved imaging from the dendrites of adult-born granule cells suggests that the cell-type specific modulation may involve an extrinsic drive to putative mitral cell-targeting granule cells.

## Results

The olfactory bulb integrates both feedforward sensory stimuli, as well as long-range projections from other brain areas (**Fig. 1A**). The latter input is thought to convey behavioural contextual signals to the olfactory bulb and tune activity patterns flexibly. To study how the behavioural context modulates the olfactory bulb output in olfactory decision making, we trained head-fixed mice to perform an olfactory discrimination task (**Fig. 1**). Water restricted mice were trained to associate a rewarded odour (S+ odour) with a water reward, and an unrewarded odour (S− odour) with no water delivery (**Fig. 1B**). Note that this paradigm includes a trace period, as we reasoned that an early cessation in the feedforward signal may maximise the chance of observing context-related activity patterns. The mice were first trained to discriminate between easily distinguishable odour mixtures, which comprised ethyl butyrate (EB) and methyl butyrate (MB), mixed at 80%/20% ratio versus a 20%/80% ratio for the S+ vs. S− stimuli, respectively (**Fig. 1C**). When the mice reached a criterion of 80% accuracy (3 ± 0.9 days, n = 6 mice, Fig. 1D), they were trained to discriminate between more similar odour mixtures (“Difficult discrimination task”; 60%/40% mixture of ethyl butyrate and methyl butyrate versus a 40%/60% mixture). This is a task known to engage many components of the olfactory bulb circuitry ^23^. Well-trained mice discriminated between these similar mixtures in 1.63 ± 0.53 s (**Supplementary fig. 1**), with comparable sniffing patterns for the S+ vs. S− odours (**Supplementary Fig. 2**), consistent with previous reports where similar odours and reward timing were used ^24,25^.

**Figure 1:**
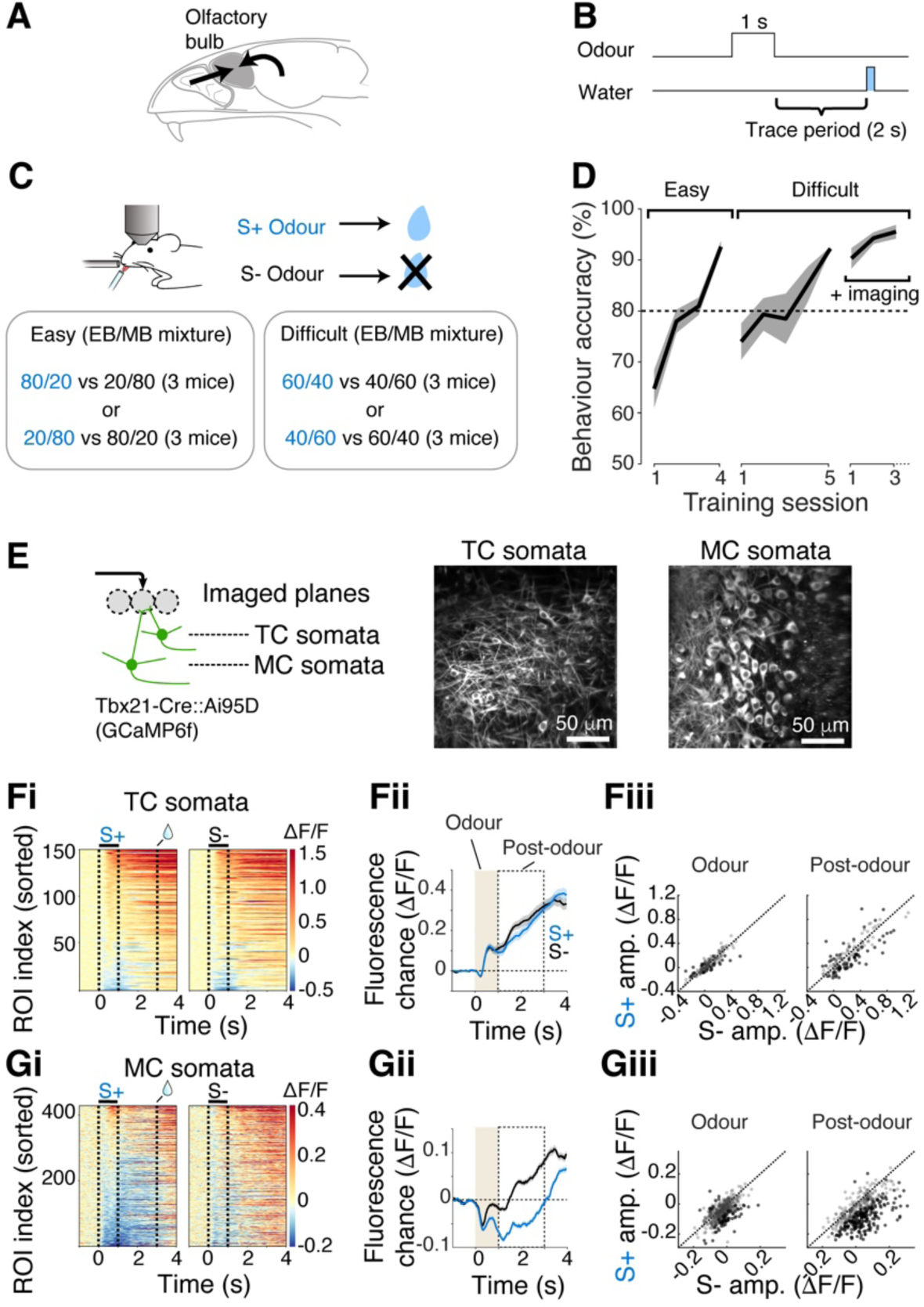
Widespread inhibition is observed in mitral cells in response to rewarded odour. **A**, Schematic showing two major sources of inputs to the olfactory bulb. Left arrow represents olfactory nerve inputs. Right arrow represents long-range inputs from other brain areas. **B**, Odour was presented for 1 s and did not overlap with the reward that was delivered 2 s after the odour offset. **C**, Odours used in the Go/NoGo olfactory discrimination tasks. Blue font corresponds to the rewarded (S+) odour. Mice were head-fixed and had a cranial window implanted. **D**, Time course of task acquisition for easy and difficult discriminations defined in **C**. Imaging sessions took place in proficient mice (n = 6 mice). **E**, Left, imaging configuration. Mitral cell (MC) and tufted cell (TC) somata were distinguished by depth. Right, example fields of view for TC somata and MC somata. **Fi-iii**, Responses to S+ and S− odours measured in TC somata. **Fi**, Colormap representation of fluorescence change over time for all ROIs. **Fii**, Average responses to S+ (blue) and S− (black) odours from all ROIs. **Fiii**, Scatter plot comparing S− vs. S+ response amplitudes for the period shown in **Fii**. Dotted line represents unity (S− amplitude = S+ amplitude). Individual points correspond to ROIs. Black dots indicate S+ and S− responses that were significantly different. **Gi-iii**, Same as **Fi-iii**, but for MC somata. N = 150 and 428 ROIs, and 3 and 6 mice for TC somata and MC somata, respectively.

In the mice proficiently performing the difficult olfactory discrimination task, we studied the responses of olfactory bulb output to the rewarded vs. unrewarded odours. The calcium indicator GCaMP6f was expressed in mitral and tufted cells using *Tbx21-Cre* mice crossed with *Ai95D* mice ^26,27^, and was imaged using a two-photon microscope (n=428 ROIs in 6 mice, and n=150 ROIs in 3 mice, respectively; **Fig. 1E-G**). Mitral and tufted cells were distinguished by depth (**Fig. 1E**). Tufted cells responded largely similarly to both odours (mean ΔF/F during odour = 0.628 ± 0.135 and 0.655 ± 0.148 for S+ and S− respectively; p = 0.777, Wilcoxon rank-sum test; mean ΔF/F post odour = 0.203 ± 0.264 and 0.237 ± 0.264 for S+ and S− respectively; p = 0.149, Wilcoxon rank-sum test; **Fig. 1F**). Peculiarly, responses of the mitral cell somata to the rewarded odour were characterised by widespread inhibitory responses (Mean ΔF/F S+ = −0.048 ± 0.058; S− = −0.022 ± 0.054; p < 0.001, Wilcoxon rank-sum test; **Fig.1G**). This dominance of inhibition for the S+ odour was present soon after the odour onset, but was particularly pronounced during the post-odour period (Mean ΔF/F S+ = −0.048 ± 0.095; S− = 0.034 ± 0.102; p < 0.001, Wilcoxon rank-sum test; **Fig. 1G**).

The late onset of the reward-associated inhibition in mitral cells raises the question regarding the underlying drive: Is the inhibitory component locked to the anticipatory motor output, or to the odour? To analyse this, we divided the rewarded trials into two sets based on the animals’ reaction times (“early onset” vs. “late onset”), and reverse-correlated the GCaMP6f signals to the onsets of anticipatory signals (“lick-aligned average”; **Fig. 2**). If the peak of inhibition in the averages occur at the same time for the early lick sets and late lick sets, it would imply the inhibition is locked more to the behavioural output (**Fig. 2B**). This analysis revealed, in contrast, that the time of peak inhibition is shifted depending on the reaction time (Pearson’s correlation coefficient = −0.555, p = 0.026; N = 15 fields of view, 6 mice; **Fig. 2C-E**), indicating that the inhibition is locked to the odour.

**Figure 2:**
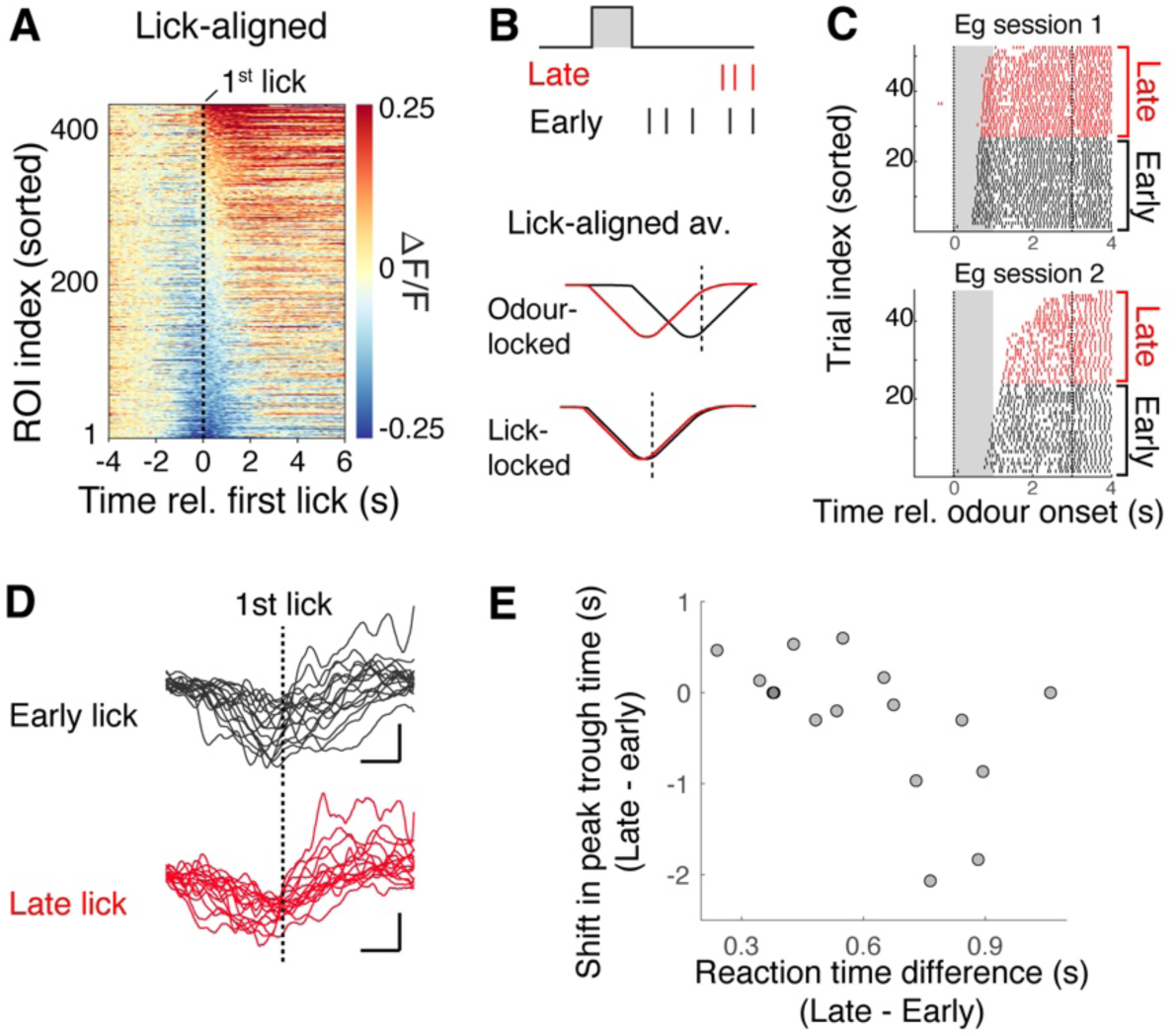
Reward-related inhibition is locked to the odour presentation. **A**, Fluorescence change from mitral cell somata aligned to the time of 1^st^ anticipatory lick after odour onset (S+ trials). **B**, Predictions for two alternative hypotheses for late-lick vs. early-lick trials; if inhibition is odour-locked, a shift in the peak inhibition is observed in the lick-aligned average. If the reward-related inhibition is locked to generation of licks, the trough times will be the same for late vs. early lick trials relative to the onset time of anticipatory lick. **C**, Lick raster plots from two example sessions. Late (black) vs. early (red) lick trials were defined as trials where first anticipatory lick occurred later or earlier than the median lick onset time for each session. **D**, Lick-aligned averages for early vs. late lick for all sessions. Each trace is an average for one session. **E**, Shift in the peak trough time in the lick-aligned average compared against mean difference in the lick onsets for the late vs. early trials. Each point corresponds to one imaging session. Pearson’s correlation coefficient = −0.555, p = 0.026 (N = 16 sessions, 6 mice).

The prevalence of inhibitory responses in mitral cells following the rewarded odour presentation is striking, but this level of inhibitory dominance has not been reported previously, even though several studies already studied how mitral cells respond to odours during difficult odour discrimination paradigms ^28–30^. The difference here may be the short duration of odour pulse used, followed by a two seconds-long trace period. It is possible that, with a longer odour presentation, the feed-forward component may dominate over any modulatory influences in the olfactory bulb (**Fig. 3A**). To test this possibility, in well-trained mice, we presented the odours for a longer period (4 s), making the task a delay task (**Fig. 3B**). In this condition, mitral cells responded to the rewarded and unrewarded odours similarly (**Fig. 3C-E**). Notably, both responses were characterised by widespread inhibitory component (% of ROIs showing significant inhibition = 18.1 for S+ and 11.9 for S−, and 35.2 for S+ and 23.9 for S− in early and late time windows, respectively, n = 5 mice). The divergent responses may therefore originate from a conjunction of olfactory and contextual signals.

**Figure 3:**
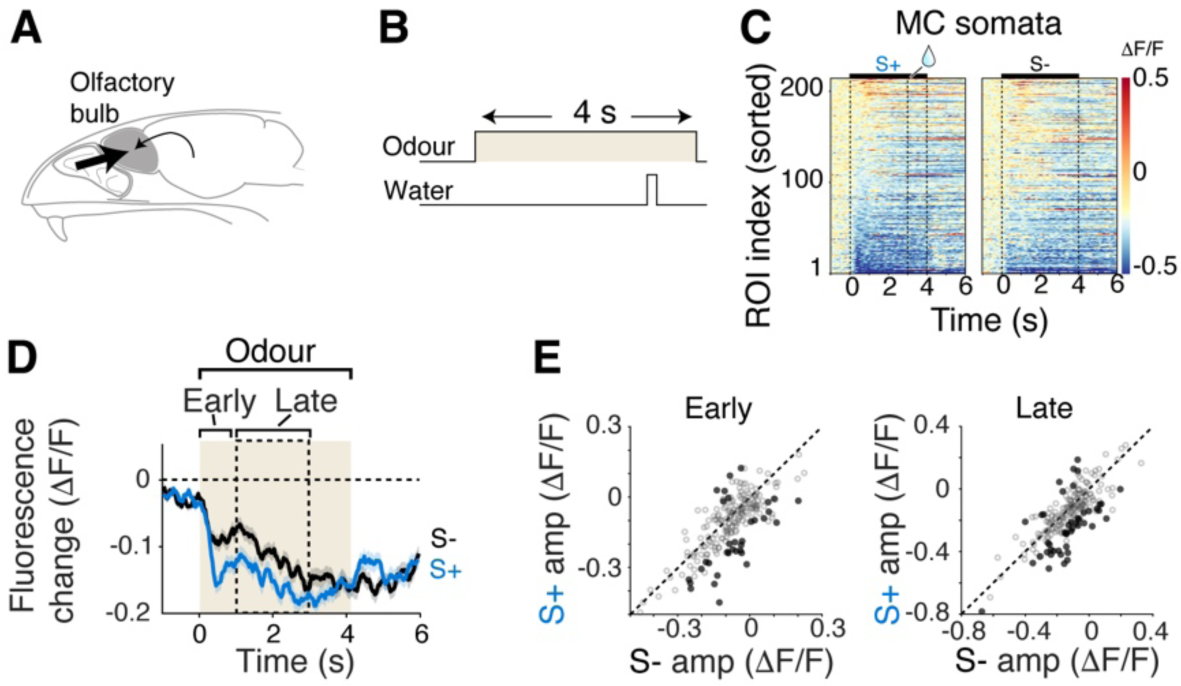
Longer odour presentation masks the appearance of divergent responses. **A**, Schematic showing dominance of sensory drive. **B**, A four-second dour pulse overlapped temporally with reward delivery. **C**, Colour map display showing GCaMP6f fluorescence change from mitral cell somata evoked by S+ and S− odours. **D**, Average fluorescence change from all ROIs in response to S+ (blue) and S− (black) odours. Mean and SEM shown. **E**, Scatter plots of average fluorescence change for S+ vs. S− odours for the time period indicated in **D**. Black dots indicate S+ and S− responses that were significantly divergent. (N = 210 ROIs, 5 mice).

To test if the behavioural state of the animal is crucial for the response divergence in mitral cells, we used two pseudo-conditioning paradigms using the same odours (**Fig. 4A,B**). In the first case (“Disengagement”), the water was delivered every trial, approximately 15 seconds before the odour presentation (**Fig. 4B**). In the second case (“Random association”), we delivered the water on randomly selected trials, so that both 60/40 and 40/60 odour mixtures were followed by water 50% of the time (**Fig. 4B**). These two paradigms decouple the odour-reward association, while inducing different levels of engagement in the head-fixed mice ^31^. Imaging sessions took place after the mice, previously trained on the fine discrimination task, were switched to, and experienced at least one session of the new paradigm (**Fig. 4C**). In both control paradigms, the head-fixed mice showed no preferential licking for the 60/40 mixture (average anticipatory licks for disengagement paradigm = 1.5 ± 1.6 and 0.7 ± 0.8 on 60/40 and 40/60,respectively; p = 0.999, 1-way ANOVA with *post-hoc* multiple-comparisons; average anticipatory licks for random association paradigm = 6.9 ± 6.1 and 5.5 ± 4.3 on 60/40 and 40/60, respectively; p = 0.890, 1-way ANOVA with *post-hoc* multiple-comparisons, **Fig. 4D**). Importantly, disengagement and randomized paradigms differed in the general levels of anticipatory licks (average anticipatory licks for all trials = 1.1 ± 1.3 and 6.2 ± 5.2 for disengagement and random association paradigms, respectively; p = 0.0021, 1-way ANOVA with *post-hoc* multiple-comparisons), indicating that different levels of behavioural engagement were indeed achieved by these paradigms. In both cases, the mitral cell somata responded similarly to the two odour mixtures (mean ΔF/F for disengagement = −0.02 ± 0.05 and −0.01 ± 0.06 during odour for 60/40 and 40/60, respectively; p = 0.465; for post odour = 0.05 ± 0.09 and 0.05 ± 0.09; p=0.553; mean ΔF/F for random association = −0.03 ± 0.09 and −0.04 ± 0.10 during odour; p = 0.259; post odour = −0.01 ± 0.14 and 7.7 x 10^−5^ ± 0.14; p=0.617, Wilcoxon rank-sum test; **Fig. 4E-G**). Note that the inhibition during the post-odour, anticipatory period that is normally present in discriminating mice was generally reduced in the two control paradigms. Together, these results indicate that the observed divergent responses in mitral cell somata are state-dependent, and not explained by the odour identities.

**Figure 4:**
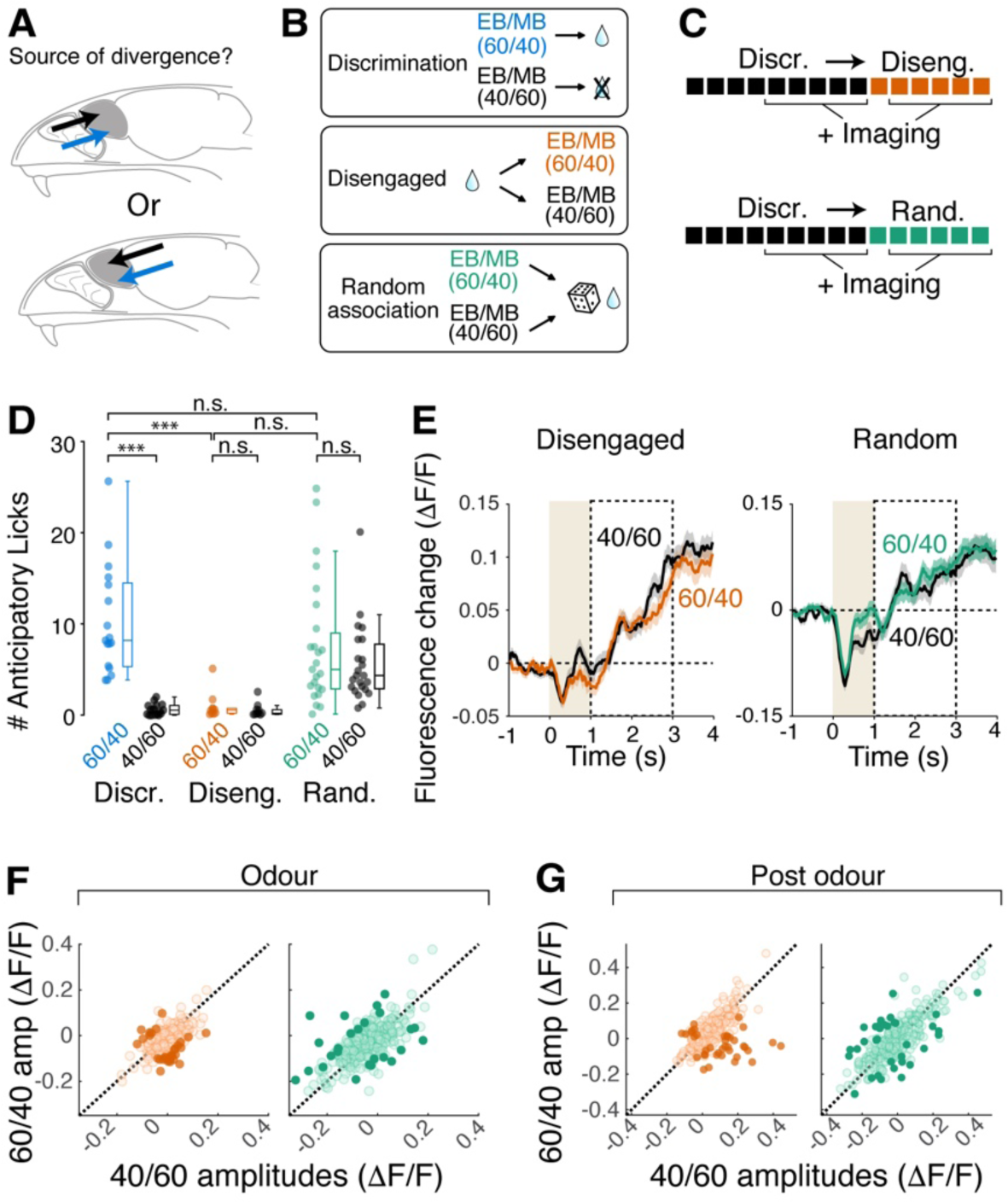
Occurrence of reward-related inhibition depends on the behavioural context, not odour identity. **A**, Hypotheses on the source of signals underlying differential S+ and S− responses in mitral cell somata; it could derive from sensory stimuli (top) or from long-range inputs to the olfactory bulb (bottom). **B**, Behavioural paradigms to decouple reward association while disengaging mice (middle) or engaging mice (Random association). In disengagement sessions, reward was delivered every trial, preceding odour presentations. In random association, reward followed both mixtures of EB and MB 50% of the time. **C**, Timeline of experiments. Mice first performed fine olfactory discrimination, then went through either disengagement or random association sessions. Imaging took place from day 2 in both cases. **D**, Number of anticipatory licks (licks 3 seconds’ window from odour onset) for the two odours for three behavioural paradigms. Individual points correspond to each imaging session analysed. *** corresponds to p < 0.001 (*post-hoc* Tukey-Kramer multiple comparisons after 1-way ANOVA) **E**, Average fluorescence change of all ROIs (mitral cell somata) for the odours indicated. **F**, Comparison of fluorescence change in response to the the two odours for the odour period for disengagement sessions (left) and random association sessions (right). Individual points correspond to ROIs. Darker points represent significantly divergent responses. **G**, Same as F, but for post-odour period. N = 125 ROIs, 3 mice for disengagement and 301 ROIs, 5 mice for random association.

What is the cellular origin of the widespread inhibition associated with the rewarded odour? Previous studies showed that a variety of feedback and neuromodulatory projections to the olfactory bulb modulate the physiology of olfactory bulb neurons ^18–20,32–34^. Further, several studies showed that such modulations manifest differently for mitral cells and tufted cells ^17,18,29,35^. Recent works indicate that mitral cells receive more potent feedback modulation from the piriform cortex ^17,18^. Thus, even though it is beyond the scope of the current work to systematically investigate all sources, the piriform cortex is a reasonable candidate for the source of the contextual signal resulting in the mitral cell-specific, reward-related inhibition we observe.

To test the involvement of the piriform cortex, we pharmacologically inactivated the ipsilateral anterior piriform cortex while the head-fixed mice performed the difficult olfactory discrimination task (**Fig. 5A**). This was achieved by infusing the GABA_A_ antagonist, muscimol, through an implanted canula. Muscimol and control sessions were carried out on alternate days, but the same fields of view were sampled for the two conditions, so that the responses of the same ROIs could be compared directly. The infusion of muscimol disrupted the behavioural performance significantly (behavioural accuracy = 64.0 ± 14.5 % during muscimol sessions; 92.8 ± 7.8 % during control sessions; p = 0.004, Wilcoxon rank-sum test; n=6 control sessions and 6 muscimol sessions, 3 mice; **Fig. 5F**). When responses of mitral cells were imaged in this condition, divergence in the rewarded vs. unrewarded odour responses were significantly reduced (Mean ΔF/F during odour = −0.007 ± 0.073 and −0.023 ± 0.091 for S+ and S− respectively; p = 0.174, Wilcoxon rank-sum test; Mean ΔF/F post-odour = 0.067 ± 0.124 and 0.079 ± 0.171 for S+ and S− respectively; p = 0.252, Wilcoxon rank-sum test, **Fig. 5G-I**). This was characterised by a reduction in the inhibitory responses evoked by the rewarded stimulus (normalised S+ ─ S− difference = −0.007 ± 0.156 and 0.043 ± 0.142 in control and muscimol sessions respectively; p = 0.008, Wilcoxon rank-sum test; **Fig. 5J**), and during the post-odour phase (normalised S+ ─ S− difference = −0.263 ± 0.175 and −0.040 ± 0.168 in control and muscimol sessions respectively; p = 3.53 x 10^−18^, Wilcoxon rank-sum test; **Fig. 5J**). Together, these data indicate that an intact piriform cortex and/or accurate behavioural performance, is required to observe the widespread inhibitory responses associated with the rewarded odour.

**Figure 5:**
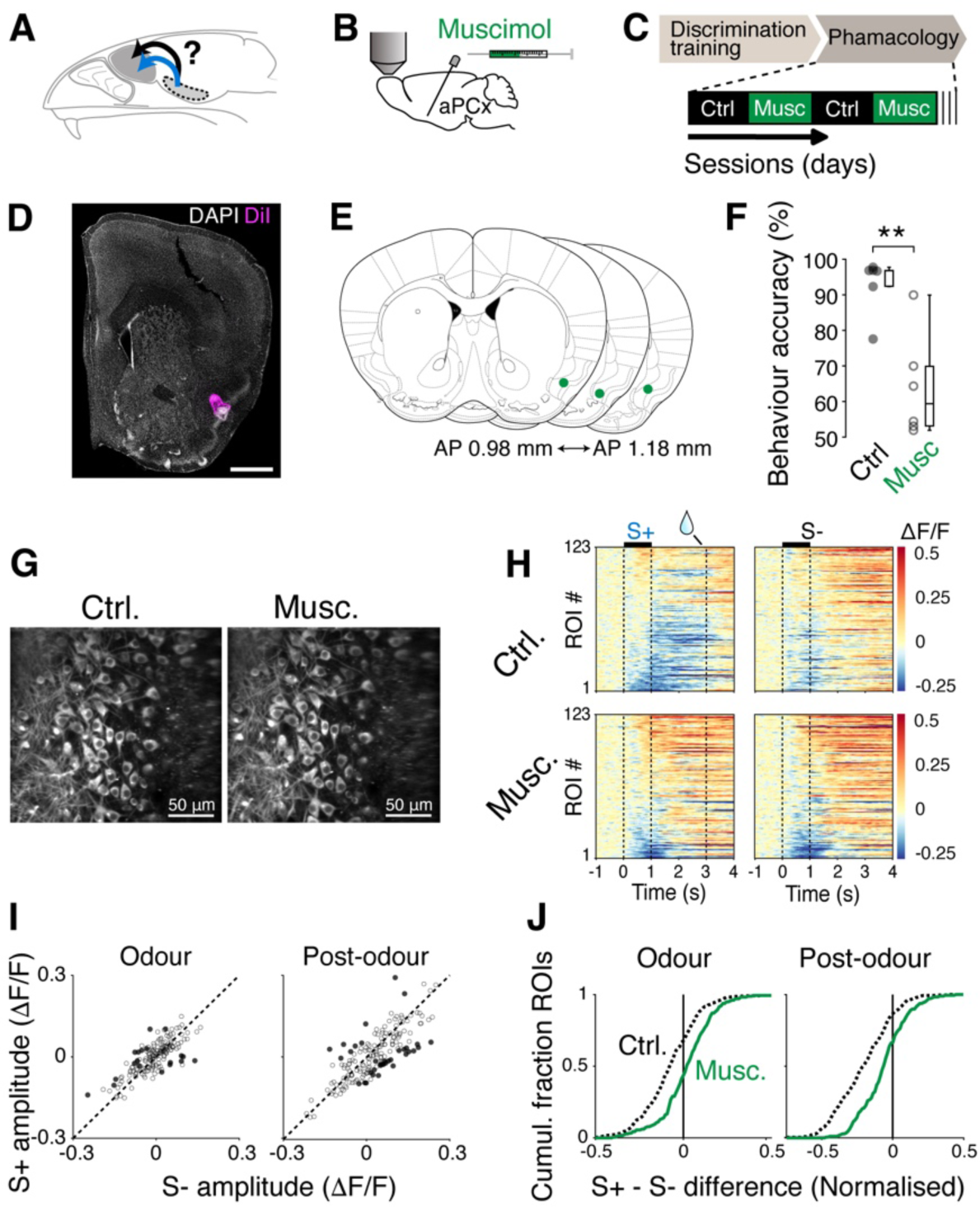
Intact piriform cortex is needed to observe the reward-related inhibition. **A**, Hypothesis tested; anterior periform cortex is necessary for mitral cell divergence during odour discrimination. **B**, Muscimol solution (2 mM; 500 nL) was infused via an implanted cannula targeted to anterior piriform cortex. **C**, Timeline of experiments. After mice were trained on the discrimination task, control and muscimol sessions alternated. One imaging session occurred per day. **D**, Example of DiI location infused via an implanted cannula. Scale bar = 1 mm. **E**, Summary of cannula tip locations. **F**, Accuracy in performance (% of trials with correct lick response) for control vs. muscimol sessions. Individual points correspond to each imaging session. P = 0.004. **G**, Example fields of view matched across two conditions. **H**, Timecourse of fluorescence change for control (top) and muscimol (bottom) sessions in matched ROIs. **I**, Average fluorescence change for each ROI for S+ and S− odours during odour and post-odour periods. Darker points represent significantly divergent responses. **J**, Cumulative fraction of ROIs for normalized difference in fluorescence changes evoked by S+ and S−. N = 124 ROIs, 3 mice.

The results so far indicate that the widespread inhibitory responses associated with the rewarded odours come from sources extrinsic to the olfactory bulb. One of the major targets of such long-range projections within the olfactory bulb is the granule cells. These cells contact mitral cells on their lateral dendrites at a deeper portion of the external plexiform layer. If the granule cells convey the contextual signals to mitral cells, the divergent responses may be observable perisomatically, but not in the superficial compartment (Fig. 6A). To test this, we compared the GCaMP6f signals from the apical dendrites of mitral cells in the glomeruli, vs. signals from the somata, which reflect signals derived from all subcellular compartments. Since tufted cells and mitral cells both send their apical dendrites to the glomeruli, to study signals from mitral cells in isolation, we used Lbhd2CreERT2::Ai95D mice, where GCaMP6f is expressed predominantly in mitral cells ^24^ (**Fig. 6A,B**).

**Figure 6:**
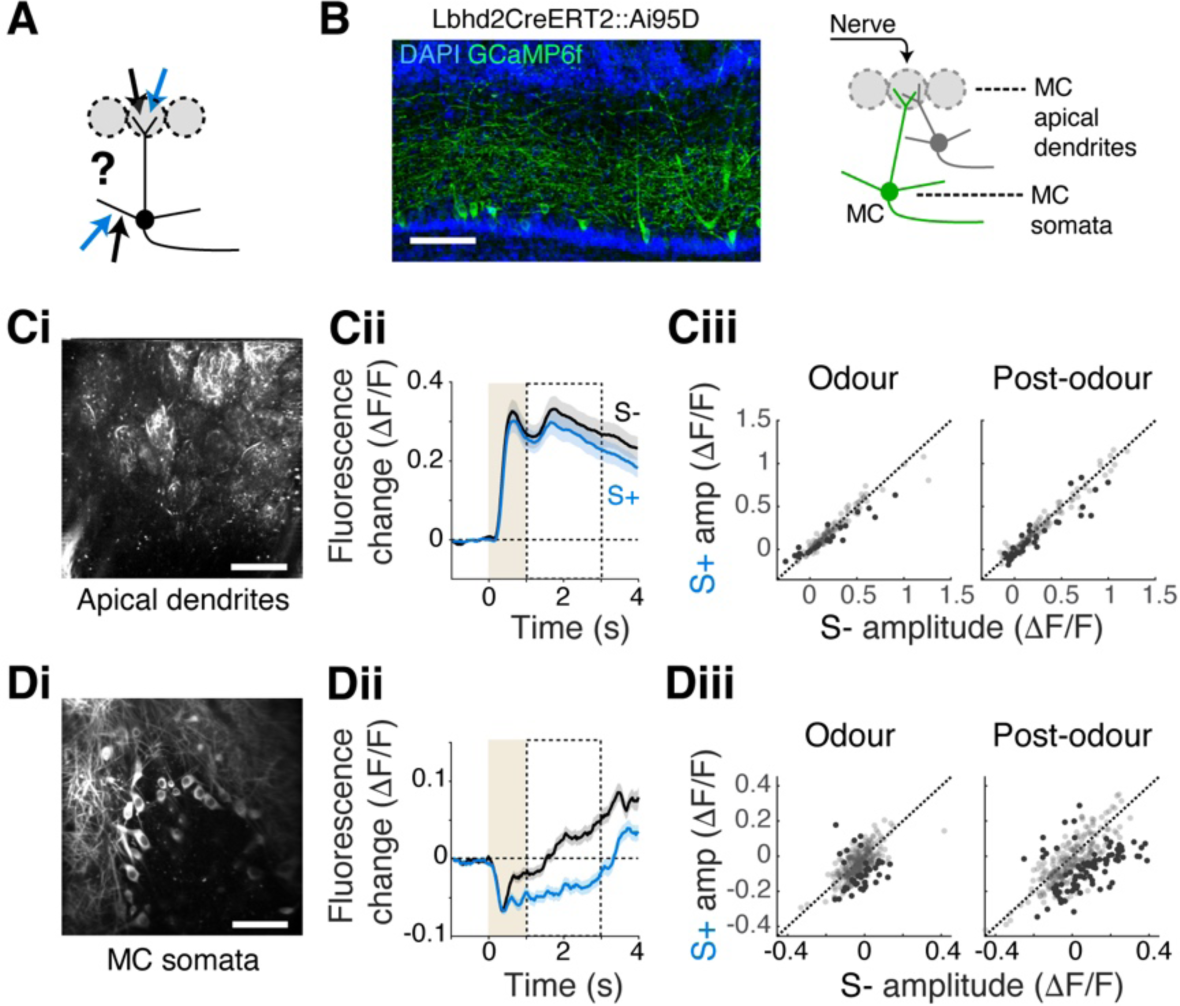
Reward-related inhibition originates peri-somatically. **A**, Schematic showing possible subcellular compartments where divergent signals may arrive, namely, apical dendrites in the glomerulus (upper arrows), and deeper, lateral dendrites (lower arrows). **B**, Left, Confocal image showing GCaMP6f preferentially in mitral cells (MCs) in Lbhd2CreERT2::Ai95D mice. Right, Illustration of imaging planes to obtain signals from the MC apical dendrites and somata. Scale bar = 100 μm. **C**, Analysis of GCaMP6f signals from MC apical dendrites. **Ci**, Example field of view from the glomerular layer. **Cii**, Average fluorescence change from all ROIs (glomeruli) in response to S+ and S− odours. **Ciii**, Comparison of average GCaMP6f fluorescence change for individual ROIs evoked by S+ vs. S− odours for the periods indicated. Darker points represent significantly divergent responses. **Di-iii**, Same as **Ci-iii**, but for MC somata. Scale bar = 50 μm. N = 140 ROIs, 4 mice for apical dendrites and 321 ROIs, 7 mice for somata.

Imaging from the superficial plane, the apical dendrites showed no significant differences between responses to S+ and S− odours (mean ΔF/F during odour = 0.307 ± 0.439 and 0.328 ± 0.466 for S+ and S− respectively; p = 0.687; post odour = 0.338 ± 0.512 and 0.382 ± 0.549 for S+ and S− respectively; p = 0.423, Wilcoxon rank-sum test, **Fig. 6C**). As before, signals from the mitral cell somata imaged in the *Lbhd2-CreERT2::Ai95D* mice were characterised by the widespread inhibitory component (mean ΔF/F during odour = −0.058 ± 0.077 and −0.034 ± 0.061 for S+ and S− respectively; p = 1.86 x 10^−4^; post-odour = −0.024 ± 0.130 and 0.038 ± 0.131 for S+ and S− respectively; p = 1.08 x 10^−5^, Wilcoxon rank-sum test; **Fig. 6D**). Together, these data suggest that the reward-related inhibition in mitral cells originates perisomatically.

If the inhibition in response to the rewarded cue in mitral cells is mediated via granule cells, we should observe a greater GCaMP6f signal change to S+ odours specifically in the granule cells that target mitral cells (**Fig. 7A**). The granule cells whose dendrites ramify in the deeper portion of the external plexiform layer are thought to synapse with mitral cells ^36–38^, where mitral cell lateral dendrites are found. These mitral cell-targeting granule cells are, however, present intermixed with tufted cell-targeting granule cells.

**Figure 7:**
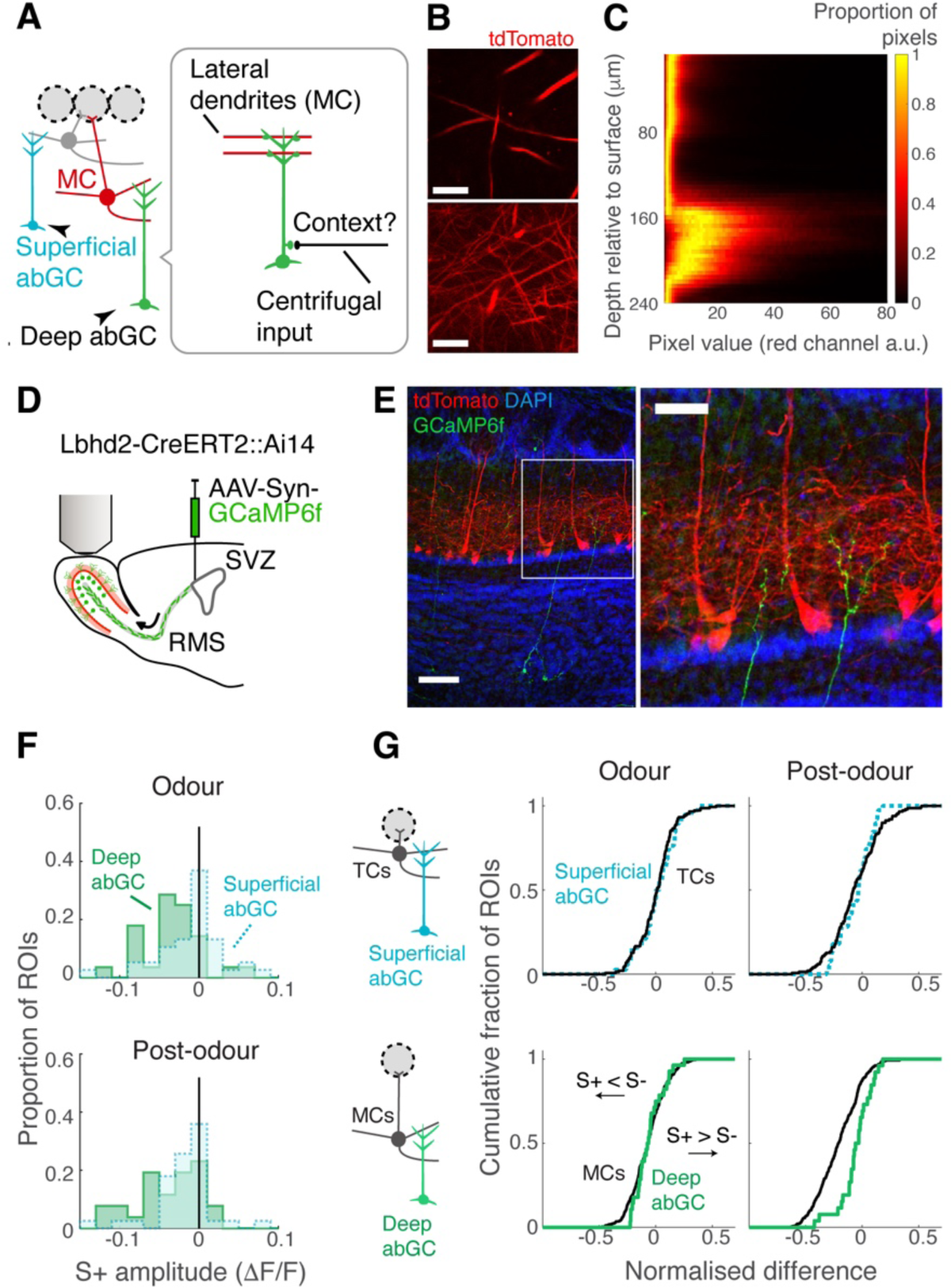
Imaging from deep adult-born granule cells suggests an extrinsic drive for mitral-targeting granule cells. **A**, Schematic of local circuitry and hypothesis; mitral cells (MCs) synapse with granule cells (GCs) whose dendrites ramify in the lower portion of EPL. The deep-ramifying granule cells may receive the contextual signal that leads to the divergent responses in MC somata. **B**, Lower and upper portions of external plexiform layer can be distinguished by the density of MC dendrites. tdTomato is preferentially expressed in MCs in Lbhd2CreERT2::Ai14 mice. Scale bars = 30 μm for top and bottom images. **C**, Depth-dependent pixel intensity histogram from an example z-stack obtained with a two-photon microscope in a Lbhd2CreERT2::Ai14 mouse. **D**, Adult-born granule cells (abGCs) are made to express GCaMP6f by injecting AAVs in the subventricular zone (SVZ). Adult-born granule cells are imaged 4 weeks after injection. RMS = rostral migratory stream. **E**, Example confocal images showing amplified GCaMP6f signal from deep abGC (green), shown with tdTomato signals (red) from MCs and DAPI signals (blue). Scale bars = 100 μm and 50 μm for left and right images, respectively. **F**, Comparison of S+ vs. S− response difference distributions for deep abGCs (green) and superficially ramifying abGCs (light blue, dotted line). **G**, Left, Comparison of S+ vs. S− response difference distributions for tufted cell (TC) somata (black trace) and superficially ramifying abGCs (light blue); right, Comparison of S+ vs. S− response difference distributions for MC somata (black trace) and deep abGCs (green).

To distinguish the putative mitral cell-targeting granule cells from those that target tufted cells, the depth of external plexiform layer needs to be distinguished accurately *in vivo*. Towards this end, we crossed *Lbhd2-CreERT2* mice with *Ai14* mice to express tdTomato preferentially in mitral cells. We reasoned that despite tissue curvature or non-uniform thickness of the external plexiform layer, this method would allow us to accurately separate the deeper from the superficial portions based on the density and distribution of the tdTomato expression. Indeed, the deep portion of the external plexiform layer showed higher density of thin red fluorescent processes (**Fig. 7B,C**), while at more superficial depths, we observed occasional fluorescence from thick processes, likely corresponding to the primary dendrites of mitral cells.

To study if the divergent odour responses in mitral cells can be explained by the evoked activity of putative mitral cell-targeting granule cells, we turned to adult-born granule cells that develop their dendrites in the deep external plexiform layer (**Fig. 7D,E**). Adult-born granule cells are thought to be critical for refining odour responses in mitral and tufted cells when mice need to discriminate between similar odours ^39–43^. Further, since the mature adult-born granule cells form dendro-dendritic synapses with mitral cell lateral dendrites, where GABA release can occur locally ^44^, we sought to image directly from dendritic gemmules. Due to their small size, we were cautious to exclude images from sessions that showed motion artefact, which was determined by correlating the structural fluorescence pattern to the baseline period and discarded those that showed low correlation (**Supplementary Fig. 3**). As a result, 70% (1343/1917 trials) of the acquired data was discarded. Deep dendritic gemmules showed more frequent inhibition compared to superficially located dendritic gemmules (**Fig 7F, Supplementary Fig. 4**). This may reflect the reduced excitatory drive in the deep granule cells, due to the prevalent reward-related inhibition in mitral cells. Thus, we analysed how the evoked amplitude distribution in the deep vs. superficial dendritic gemmules relative to that of the presynaptic counterparts, that is, against the distribution of evoked responses in mitral cells and tufted cells, respectively. The S+ vs. S− tuning showed a close overlap between tufted cells and superficial gemmules of adult-born granule cells. On the other hand, S+ vs. S− tuning distribution of deep gemmules could not be explained by the mitral cell tuning distribution (**Fig. 7G**). There was a tendency for these gemmules to respond more positively to the rewarded odours than would be predicted from mitral cell activity alone. In other words, our data suggests that mitral cell-targeting granule cells may receive an additional excitatory drive associated with the rewarded odour.

## Discussion

In this study, we observed a cell-type specific reward-associated inhibition in the primary olfactory area of the mouse. This inhibition is cell-type specific and subcellular specific: first, it manifests in the somata of mitral cells but not in tufted cells, and second, it appears in the somata but not in their apical dendritic tuft in the input layer. This subcellular specificity suggests that the generation of this phenomenon involves circuit components at a deeper layer of the olfactory bulb. Further, the results of pseudo-conditioning and pharmacological manipulations suggest that the mitral cell-specific, reward-related inhibition arises from an acquired congruence of sensory and contextual signals. Thus, our study supports the notion that value-related modulation of olfactory signals is a characteristic of olfactory processing in the primary olfactory area and provides constraints on possible underlying mechanisms.

Many questions remain regarding the origin of the reward-related signals to the olfactory bulb. Many brain regions send long-range projections to the olfactory bulb and are, therefore, candidate drivers of the reward-related inhibition we observed. One important source of reward-related signals to OB is the direct or indirect feedback projections from olfactory cortices. Value-like modulation of olfactory responses occurs in many parts of the brain: It has been observed in the prefrontal cortex ^4,45^, orbitofrontal cortex ^4,45^, hippocampus ^46^, olfactory tubercle ^47–50^, piriform cortex ^4,51^, and anterior olfactory nucleus ^4^, although there may be regional differences, for example, in the long-term stability of expression ^45^. Of particular interest is the piriform cortex, which serves as a gateway for processed signals for modulation of the mitral cells ^17^. While only a subset of pyramidal neurons from the piriform cortex project to the olfactory bulb ^52^, a recent imaging study from olfactory bulb-projecting fibres showed value-like activity when the task depended on olfactory cues ^53^. It is unclear why we did not observe the widespread reward-related modulation in tufted cells, even though some value-like activity is present in the anterior olfactory nucleus ^4^, a region known to have modulatory influence over tufted cells ^17^. Since the anterior olfactory nucleus has multiple compartments ^54^ with each with distinct long-range connectivity ^55^, it will be crucial for future studies to resolve how these subregions contribute to associating values with olfactory stimuli.

Another potential source of reward-related signals are classical neuromodulatory regions. Of particular interest are those that innervate the deeper layers of the olfactory bulb, including cholinergic neurons of the nucleus of the horizontal limb of the diagonal band ^19,56^, noradrenergic neurons of the locus coeruleus ^57,58^ and serotonergic neurons of the dorsal raphe nucleus ^59,60^. These regions contain neurons that show reward-related signals, although they differ in the details. Cholinergic neurons of the basal forebrain are locked to the time of anticipatory behaviour in reward-driven sensory tasks ^61,62^. This timing is relatively late compared to our phenomenon. In addition, an optogenetic activation of cholinergic fibers in OB enhances, rather than inhibits, odour-evoked responses in mitral and tufted cells ^19^. Normal adrenergic transmission within the olfactory bulb is required for learning to discriminate similar odours ^63,64^. Neurons of the locus coeruleus, too, show instances of reward-locked activations ^65^. As with acetylcholine, however, the timing is locked more to the anticipatory actions and further, modulation may enhance, rather than inhibit, olfactory responses in the bulb during delay conditioning ^22^. Therefore, cholinergic and adrenergic inputs may not relate directly to the phenomenon described in this study. On the other hand, among serotonergic neurons of the dorsal raphe nucleus, some lock to rewarded cues ^66–68^, making the serotonergic input a promising candidate for the reward-related inhibition in mitral cells. Precisely what the serotonergic signal represents remains an active area of research, with possibilities including reward ^67^, uncertainty ^68^, or a motor-sensory variable ^69^, though this may reflect regional differences, too ^68,70^. Since serotonergic neurons also target the striatum to affect the dopamine release in the region ^71^, future experiments must resolve if they pose direct reward-related effects on OB or via other brain areas.

The value-like activity in the higher olfactory areas mentioned above appears as excitatory responses to the rewarded odour. In contrast, mitral cells of the olfactory bulb showed enhanced inhibitory responses. This is reminiscent of a human study, where perceived unpleasantness (negative valence) positively modulated the late component of odour-evoked responses in the olfactory bulb ^72^. The sign reversal in the olfactory bulb suggests an involvement of inhibitory interneurons like the granule cells. Due to the perisomatic nature of the reward-related inhibition, the granule cells are a prime candidate. They are the major recipients of long-range projections in the olfactory bulb and may target mitral cells vs. tufted cells separately by ramifying dendrites at the deep vs. superficial levels of the external plexiform layer ^36–38^. Despite their numerical dominance ^73^, how granule cells contribute to signal transformation in the olfactory bulb remains enigmatic. They are thought to contribute to the temporal precision of the OB output ^74–76^, which is perceptible to the animals ^77^. Granule cells seem to contribute only subtly to mitral and tufted cells’ spontaneous and evoked firing rates under anaesthesia or when mice are not engaged in behavioural tasks ^76^. However, they can potently silence mitral and tufted cells when activated in bulk ^76^. As a result, it has been hypothesised that an excitatory drive from long-range projections onto granule cells is critical for potent physiological effects ^78^. Because feedback projections synapse more proximally to the soma, these have more significant electrical impacts at the soma than inputs from the mitral and tufted lateral dendrites arriving at the distal dendrites. This drive may be critical to a global activation of these neurons ^78^, serving as a potential associative mechanism ^44^. Our depth-specific imaging approach may open new ways to investigate the physiology of these enigmatic interneurons.

Feedback signals from higher sensory regions may allow refined or processed information to be integrated into more peripheral processing, as in the olfactory bulb. Such a system may be used to dynamically tune the nature of early sensory processing to the behavioural demands at hand. But given the inhibitory timing we observed, where the modulation occurs after decisions have already been made, feedback signals may be used to fine-tune the process of maturation of adult-born granule cells and their integration into the existing circuitry within the olfactory bulb in a behaviourally relevant manner ^39–41^. It will be an intriguing future investigation to test if disruption of feedback signals prevents the proper establishment of acquired connectivity and, as a result, the expression of task-dependent activity patterns. In summary, our work brings us closer to a mechanistic understanding of context-dependent modulation involving a congruent interaction between local and long-range inputs.

## Acknowledgement

We thank Yu-Pei Huang for technical assistance, Adam Mago and Sourjya Nath Baibhabee for help with tamoxifen administration, the dedicated staff from the Animal Resources Section, Imaging section, and mechanical engineering section of OIST Graduate University for the support, and the members of Sensory and Behavioural Neuroscience Unit for helpful comments on the project. This work was supported by Grant-in-Aid for Scientific Research (C) 22K06490 and OIST Graduate University.

## Materials and Methods

### Animals

All animal experiments had been approved by the Okinawa Institute of Science and Technology Graduate University Graduate Animal Care and Use Committee (Protocol 2020–310). *Tbx21-Cre* (Haddad et al., 2013), B6J.Cg-Gt(ROSA)26Sortm95.1(CAG-GCaMP6f)Hze/MwarJ, also known as *Ai95D* (Madisen et al., 2015), and *Ai14* (Madisen et al., 2010) mice were originally obtained from the Jackson Laboratory (stock numbers: 024507, 028865 and 007914, respectfully). *LBHD2-CreERT2* mice were generated previously and are also available through Jackson Laboratory (stock number 036054; ^24^). *Tbx21-Cre::Ai95D* and *LBHD2-CreERT2::Ai95D* mice were generated by crossing parents homozygous for each transgene. *LBHD2-CreERT2::Ai14* mice were generated by crossing LBHD2-CreERT2 mice with Ai14 mice.

### Tamoxifen administration

To induce Cre-recombinase activity in *LBHD2-CreERT2::Ai95D* and *LBHD2-CreERT2::Ai14* mice, tamoxifen injections (3 x 80 mg/kg at p21, s.c.), or tamoxifen diet (553.8 ± 362.6 mk/kg at p21) was used. For injections, each of three consecutive days, tamoxifen solution (8 mg/ml; Sigma-Aldrich T5648) was first dissolved in absolute ethanol. This was then added to corn oil, resulting in a 5% ethanol and 95% corn oil mixture, and heated at 65 °C on a shaker for 30 minutes. After it cooled down to room temperature, the solution was injected intraperitoneally (∼ 100 μl). Mice that were treated with the tamoxifen diet (2 mg/kg) were exposed to this for 2-4 days, based on their initial weight, after which they were switched back to normal food (threshold to switch to normal food: 80% initial body weight). Tamoxifen intake was calculated based on the amount of diet food provided and amount of diet food left after switching back to normal food.

### Surgery

All recovery surgeries were conducted in an aseptic manner. For the cranial window and headplate implantations, 9-11 week-old male mice were deeply anesthetised with isoflurane (3-5% for induction, 1-2% for maintenance; IsoFlo, Zoetis Japan). A craniotomy of approximately 1.5 x 1 mm was performed over the left olfactory bulb, and a custom cut glass window (thickness No. 1; Matsunami, Japan) was implanted. Once the window was sealed with cyanoacrylate (Histoacryl, B. Braun, Germany), a custom-made metal headplate (26 x 12 mm) was implanted posterior to the cranial window. Dental acrylic (Kulzer, Hanau, Germany) was then added to cover the exposed skull and to secure both the headplate and cranial window.

For experiments involving pharmacological infusion, an additional cannula (10 mm length; C315GS-4/SPC, Plastics One) was inserted to target the left anterior piriform cortex (coordinate: AP −2.2 mm & ML −2.4 mm from bregma; DV −6.1 mm at 45-degree angle from brain surface), similar to ^18^.

All mice were administered carprofen (5 mg/kg, i.p.) post-operatively for three consecutive days. All mice were recovered for at least two weeks before the start of behavioural experiments.

### Virus injection

To express GCaMP6f in adult-born granule cells, during cranial window and headplate implantations, *Lbhd2-CreERT2::Ai14* mice were injected with AAV1-syn-GCaMP6f-WPRE-SV40 (Addgene 100837-AAV1; titre was 1.84 x 10^13^ GC/mL at the time of synthesis) in the left SVZ (200 μL; coordinate: AP −1.0 mm & ML −1.0 mm from bregma; DV −2.2 mm vertically) using Nanoject III (Drummond, 3-000-207).

### Habituation

Male mice were habituated to head fixation on a custom-made running wheel. Thereafter, water access was restricted by removing water bottles from their home cages. The mice were habituated to receive water from the port at the experimental setup on the following two to three days mice, until they learned to drink at least 1 ml at the setup. The body weight was recorded daily, to ensure that it stayed above 80% of the original weight. Lick responses were measured using an IR beam sensor (PM-F25, Panasonic, Osaka, Japan).

During calcium imaging sessions, respiration of the mice was recorded using a flow sensor (AWM3100V, Honeywell, NC) placed closely to the right nostril.

### Discrimination training

After habituation, the mice were trained to associate one odour stimulus with a water reward (S+ odour) and another odour stimulus with no reward (S− odour). Both S+ and S− odours were a binary mix of ethyl butyrate (Sigma-Aldrich; W242705) and methyl butyrate (Tokyo Chemical Industry; B0763), but mixed with different ratios based on the photoionization detector readings. For the initial, easy, discrimination training, a 80/20 vs 20/80 ratio was used. When mice reached 80% behavioural accuracy, fine discrimination training started, using 60/40 vs 40/60 odour mixtures. The correct response to S+ odours was to lick within a 3-second window after stimulus onset, while the correct response to S− odours was to refrain from licking. The water reward consisted of multiple drops, with a total of ∼18 μL per trial. Olfactory stimuli were presented using a custom flow-dilution olfactometer ^79^. On each trial, odour was presented for 1 second and delay to the reward was 3 seconds from the odour onset. Inter-trial interval was approximately 20 seconds.

### Long odour discrimination

The mice proficient at the difficult discriminaition task were trained to discriminate between the same odour mixtures but with 4 seconds of odour duration. The response window and reward timing on S+ trials was the same between the two paradigms.

### Disengagement paradigm

In this paradigm, the same odour mixtures as the difficult discrimination were used, but the water reward was delivered every trial, approximately 15 seconds before the odour onset. The time window used for measuring the anticipatory licks was identical to that of the fine discrimination paradigm. The first session was considered a transition session and excluded from imaging analysis.

### Random association paradigm

In this paradigm, the water reward was presented in 50% of the trials, regardless of the odour identity. This decoupled the odour identity and reward, but kept mice engaged, indicated by the anticipatory licks. The first session was considered a transition session and excluded from imaging analysis.

### Pharmacological inactivation of anterior piriform cortex

For pharmacological inactivation of the anterior piriform cortex, muscimol (M1523, Sigma-Aldrich, MO) was infused (2 mM in Ringer; 500 nL at 100 nL/min) through the previously implanted cannula, using a Hamilton syringe (1 μL Model 7001KH PST-3 80100, Hamilton Company, Nevada USA), approximately 10 minutes before the start of the imaging sessions. Two days prior to the first muscimol infusion, Ringer solution (500 nL at 100 nl/min) was infused.

### Histology

After the conclusion of the behavioural experiments, the mice were perfused transcardially with phosphate buffer (in mM): NaH2 PO4 (225.7), Na2HPO4 (774.0) with pH adjusted to 7.4, followed by PFA solution (4% dissolved in phosphate buffer). For mice that were implanted with a cannula, 500 nL DiI (Invitrogen, V22885) was injected prior to the perfusion to mark the cannula tip location. Coronal sections of 100 µm thickness were cut on a vibratome (5100 mz-Plus, Campden Instruments, Leicestershire, UK) and counterstained using DAPI (D9542, Sigma-Aldrich). Images were acquired using a Leica SP8 confocal microscope using a ×10 (NA 0.40 Plan-Apochromat, 506407, Leica) objective.

### Immunohistochemistry

Free floating olfactory bulb sections from above were first blocked in blocking solution (0.025 M Tris-HCl, 0.5 M NaCl, 0.2% triton X-100, 7.5% normal goat serum, 2.5% BSA, pH = 7.5) for 60min at room-temperature. Sliced were subsequently stained with chicken anti-GFP (Abcam, ab13970; 1:500 in blocking solution) at 4°C over-night. Slices were washed three times in TBS (0.025 M Tris-HCl, 0.5 M NaCl, pH = 7.5) and incubated in goat anti-chicken Alexa-488 (Abcam, ab150169; 1:1000 in TBS supplemented with 0.2% triton X-100) for 2 hours at room-temperature. All slices were counter stained with DAPI (D9542, Sigma-Aldrich). Images were acquired using a Leica SP8 confocal microscope using a 10X (NA 0.40 Plan-Apochromat) or a 40X (NA 1.10 Plan-Apochromat, 506357, Leica) objective.

### In vivo calcium imaging

All the calcium data presented in this manuscript were obtained from awake mice. Two-photon fluorescence of GCaMP6f and tdTomato were measured simultaneously with a custom-made miscroscope (INSS, UK) fitted with a 25x objective (Nikon N25X-APO-MP1300, 1.1 N.A.) or a 16x objective (Nikon N16XLWD-PF, 0.8 NA), and high-power laser (930 nm; Insight DeepSee, MaiTai HP, Spectra-Physics, USA) at depths 50–400 μm below the surface of the olfactory bulb. Images from a single plane were obtained at ∼30 Hz with a resonant scanner. In each trial, 400 image frames were acquired, with 100 frames before odour stimulus to obtain a baseline. Each day, the stage coordinates were chosen relative to a reference location, which was determined by the surface blood vessel pattern. Fields of view were 512 μm x 512 μm for apical dendrites, 256 μm x 256 μm for tufted and mitral cell somata, 128 μm x 128 μm and for adult-born granule cell gemmules. Calcium data during difficult discrimination and disengaged experiments were obtained from *Tbx21-Cre::Ai95D* mice (Figs. 1 and 4). 6 male mice were used for somata imaging. All 6 were used to image mitral cell somata, while a subset (3 mice) were used to image tufted cell somata. To obtain calcium data from different subcelluar compartments of mitral cells, we used *LBHD2-CreERT2::Ai95D* mice (Fig. 6). For long odour discrimination, random association, and muscimol infusion experiments, calcium data was obtained from both *Tbx21-Cre::Ai95D* and *LBHD2-CreERT2::Ai95D* mice (Figs. 3-5). Finally, *LBHD2-CreERT2::Ai14* mice were used in to record red (tdTomato, mitral cells) and green (calcium indicator, gemmules) fluorescent signals during adult-born granule cell imaging experiments (Fig. 7).

### Data analysis

All data was analyzed offline using custom MATLAB (MathWorks, USA) routines. To calculate the behavioural accuracy, the number of licks during a 3 second window from final valve opening until reward presentation was counted for each trial (anticipatory licks). Correct response to the rewarded odour was a minimum of 2 anticipatory licks, and correct response to the unrewarded odour was less then 2 anticipatory licks. Behavioural accuracy was calculated as the percentage of correct trials from the total number of trials.

To calculate the sniffing frequency and speed of inhalation, the sniffing signal was first filtered (1 Hz high-pass and 30 Hz low-pass) and normalised (z-score). Inhalation peaks were detected using the *findpeaks* MATLAB function. Sniff onsets were determined by searching back in time from each detected inhalation peak to the point where the signal crossed a threshold value. The detected onsets and peaks were then used to calculate the frequency (as 1/inter-onset time) and speed of inhalation (as onset-to-peak time).

### Image analysis

For each field of view, the imaging data was manually curated based on motion artifacts and drift over time. Data with motion artifacts and/or drift were motion corrected using the NoRMCorre toolbox (Pnevmatikakis and Giovannucci, 2017) and, when unsuccessfully corrected, excluded from analysis. Regions of interest (ROIs) were manually drawn using ImageJ (NIH, Bethesda, USA) based on the average field of view from each imaging session and exported for usage in MATLAB. Average pixel value from each ROI was offset with a value from the darkest region in the frame (e.g. a blood vessel). To account for bleaching over the course of the imaging session, the mean pixel values for all trials were concatenated and detrended using the MATLAB function *detrend*, then reshaped back into an array (individual trials x frames) before relative fluorescence change was obtained. For each ROI, the change in fluorescence (ΔF/F) was calculated by subtracting the mean pixel value from the baseline period (1 second before odour stimulus onset), and dividing by the baseline value. Odour evoked responses were calculated as the mean fluorescence change during the odour stimulus presentation, and post-odour evoked responses as the mean fluorescence change between odour stimulus offset and reward presentation. For the ‘long odour’ and ‘random association’ experiments, the time windows to calculate the evoked responses were based on the fine odour discrimination experiments.

### Lick-aligned average

Rewarded trials were analysed for each imaging session. Onsets of anticipatory licks were defined as the average time of first two licks observed after the start of odour presentation. Rewarded trials were grouped into early vs late lick trials if the anticipatory lick onsets occurred before or after the median onset time, respectively. Within each group, calcium transients were aligned to the anticipatory lick onset time for each rewarded trial, and averaged.

### Quality assessment abGC imaging

Small abGC gemmules makes them susceptible to motion artifacts in behaving animals. To objectively assess the quality of the imaged trial, the tdTomato signals from the MC dendrites in *LBHD2-CreERT2::Ai14* mice were analysed. Rolling averages of 5 frames (step size: 1 frame) were made and compared against the average of 50 frames obtained during the baseline period to compute the correlation coefficient. If the mean correlation value for the period analysed (between the odour offset and onset of water reward) was below 95% of the mean value during the baseline period, the trial was rejected. Separate quality check was conducted for the odour and post-odour phase. Further, trials where the baseline correlations deviated significantly were considered outliers and rejected. This was assessed using the Matlab function *isoutlier*. This quality control method resulted in 16 accepted fields of view from 5 mice for the odour period, and 13 fields of view from 4 mice for the post-odour period. On average, a given accepted fields of view yielded 3.2 ± 1.4 ROIs (3.8 ± 1.5 ROIs for deep FOVs, and 2.9 ± 1.4 ROIs for superficial fields of view for the magnification used (128 μm x 128 μm frame size).

### External plexiform layer depth determination based on red fluorescence

Depth within the external plexiform layer was estimated using the red fluorescence signal from mitral dendrites in Lbhd2-CreERT2::Ai14 mice, which is dense in the deeper portion. Z-stack ranging from the superficial layer to mitral cell layer spanned 250 μm (100 frames averaged every 4 μm) obtained from the same x-y location as the functional imaging was used. Since the fibre-like structures are the relevant signals, the averaged frame from each depth was passed through a filter available as a plug-in in ImageJ (“Tubeness”; ^80^), with the sigma parameter set to 2 μm.

### Normalised difference

For each trial, the average value of relative fluorescence change was calculated for the odour period (first 1 second after the odour onset) and the post-odour period (1-3 s after the odour onset). The normalized difference between S+ and S− response amplitudes for the odour period, as well as the post-odour period, is calculated as follows:

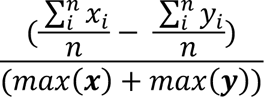

where *i* denotes the trial index, *n* is the number of trials, x is the evoked fluorescence change in response to the rewarded odour, and y is the evoked fluorescence change in response to the unrewarded odour.

### Statistics

#### Divergent responders

To determine if a ROI showed a divergent response, for each ROI odour evoked and post-odour response amplitudes for S+ and S− trials were tested for statistical significance using the Wilcoxon rank sum test. Summary transients presented in figures show mean ± SEM, unless otherwise stated.

#### Discrimination time

The method for determining the discrimination time was modified from ^81^. Cumulative histograms of the detected licks were calculated for all trials using 50 ms time bins. For each time bin, the histogram values for S+ trials and for S− trials were tested for statistical significance using a Wilcoxon rank sum test. The first time bin where histograms were significantly different (p-value less than 0.05) was taken as the discrimination time.

---

**Supplementary figure 1:**
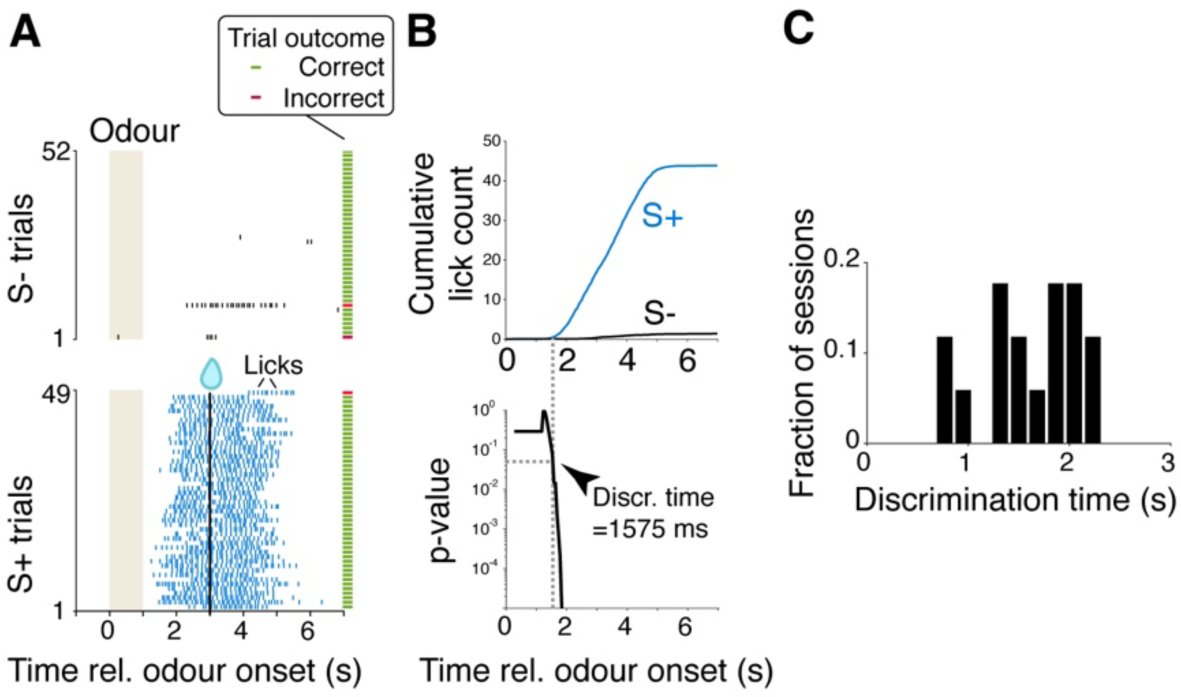
Time-course of decision-related behavioural output. **A**, Example raster plots showing lick times relative to the onset of odour (t = 0) and reward delivery (t = 3 s) of a proficient mouse. Trials have been sorted into rewarded (S+) and unrewarded (S−) trials. Whether the mice made the correct or incorrect decision was determined based on the number of licks observed between the odour onset and reward onset. Green ticks on the right indicate correct trials, and red ticks indicate incorrect trials. **B**, Calculation of discrimination time is the earliest time at which licking patterns for S+ and S− trials diverge significantly, at the 0.05 level, shown for the example session in **A**. **C**, Discrimination time for all mice presented in Figure 1. N = 17 sessions, 6 mice.

**Supplementary figure 2:**
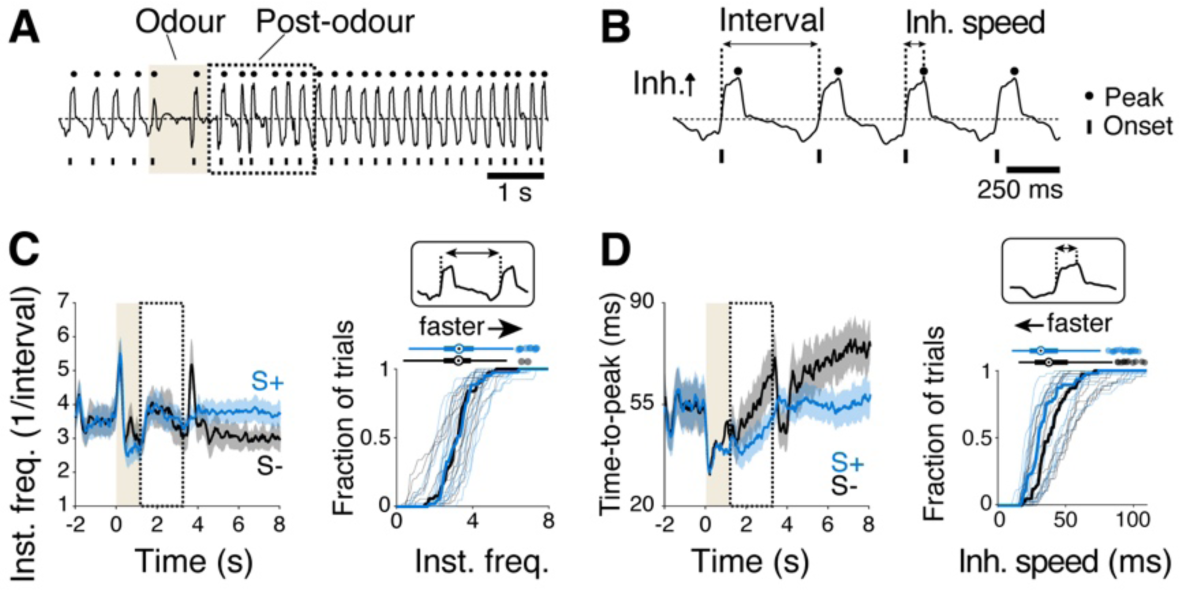
Sniff patterns are not starkly different between rewarded vs. unrewarded trials. **A**, Example sniff signal from a flow sensor placed next to a nostril on the contralateral side to odour presentation. Upward signal (signal above dotted horizontal line) corresponds to inhalation. Inhalation peaks are shown with circles (top) and exhalation peaks are annotated with short vertical ticks (bottom). **B**, Instantaneous sniff frequency is defined as the reciprocal of the sniff interval, measured from one inhalation onset to the next inhalation onset. Speed of inhalation (time to peak) is defined as the time elapsed from the inhalation onset to the inhalation peak. **C**, Time course of instantaneous sniff frequency change relative to the odour period (light brown background) and post-odour period (demarcated with dotted lines). Mean and SEM shown. **D**, Cumulative histogram of instantaneous frequencies observed during post-odour period. Thin lines correspond to individual sessions, and thick lines correspond to average across imaging sessions. **E**, same as **D**, but for inhalation speed. N = 13 sessions, 7 mice.

**Supplementary figure 3:**
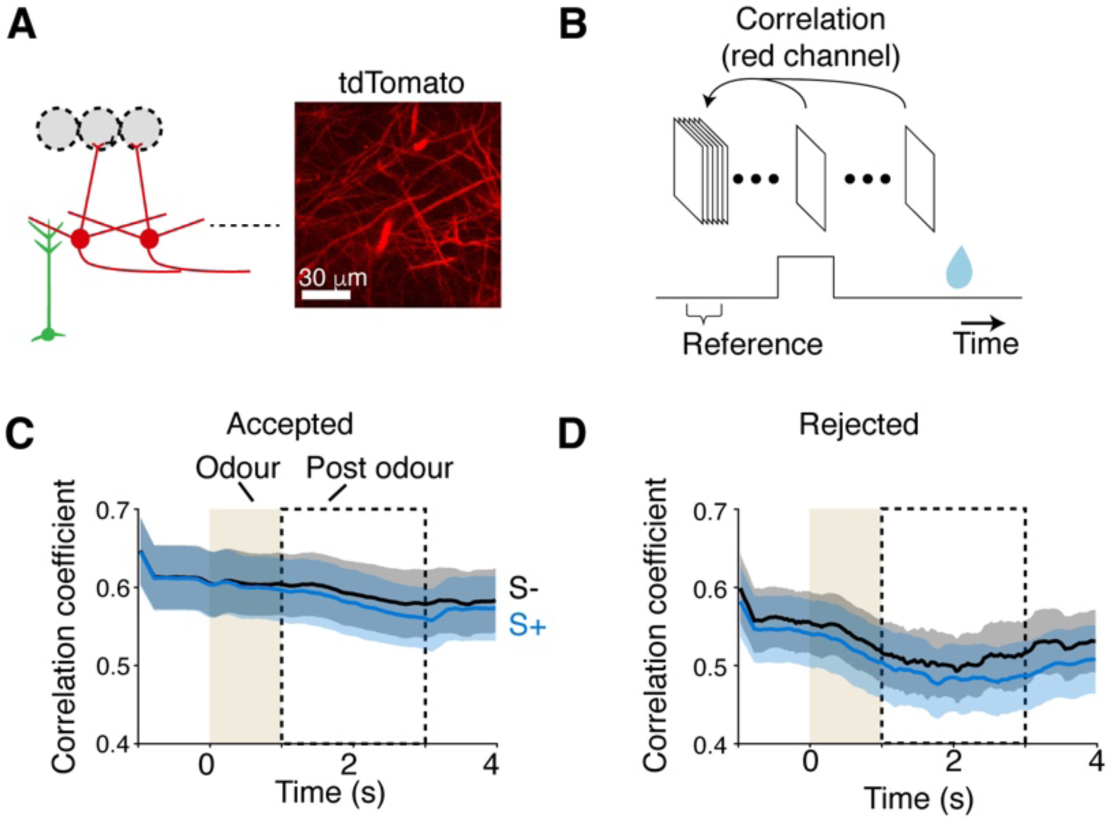
A method for assessing the quality of adult-born granule cell imaging. **A**, Example field of view from the red channel showing mitral cell dendrites. **B**, Image quality was determined by the frame-by-frame similarity of red fluorescence patterns by calculating correlation in the tdTomato image between the baseline period and other time points within the trial. Those with a high correlation coefficient throughout the trial is deemed to have less drift e.g., due to animal’s movements. **C**, Time course of red fluorescence correlation values for the accepted dataset. **D**, Same as **C** but for rejected dataset. Of the 1917 trials imaged in total 574 trials were accepted and 1343 trials were rejected. This amounts to, on average, 27.7% acceptance rate for deep gemmules and 36.2% acceptance rate for superficial gemmules.

**Supplementary figure 4:**
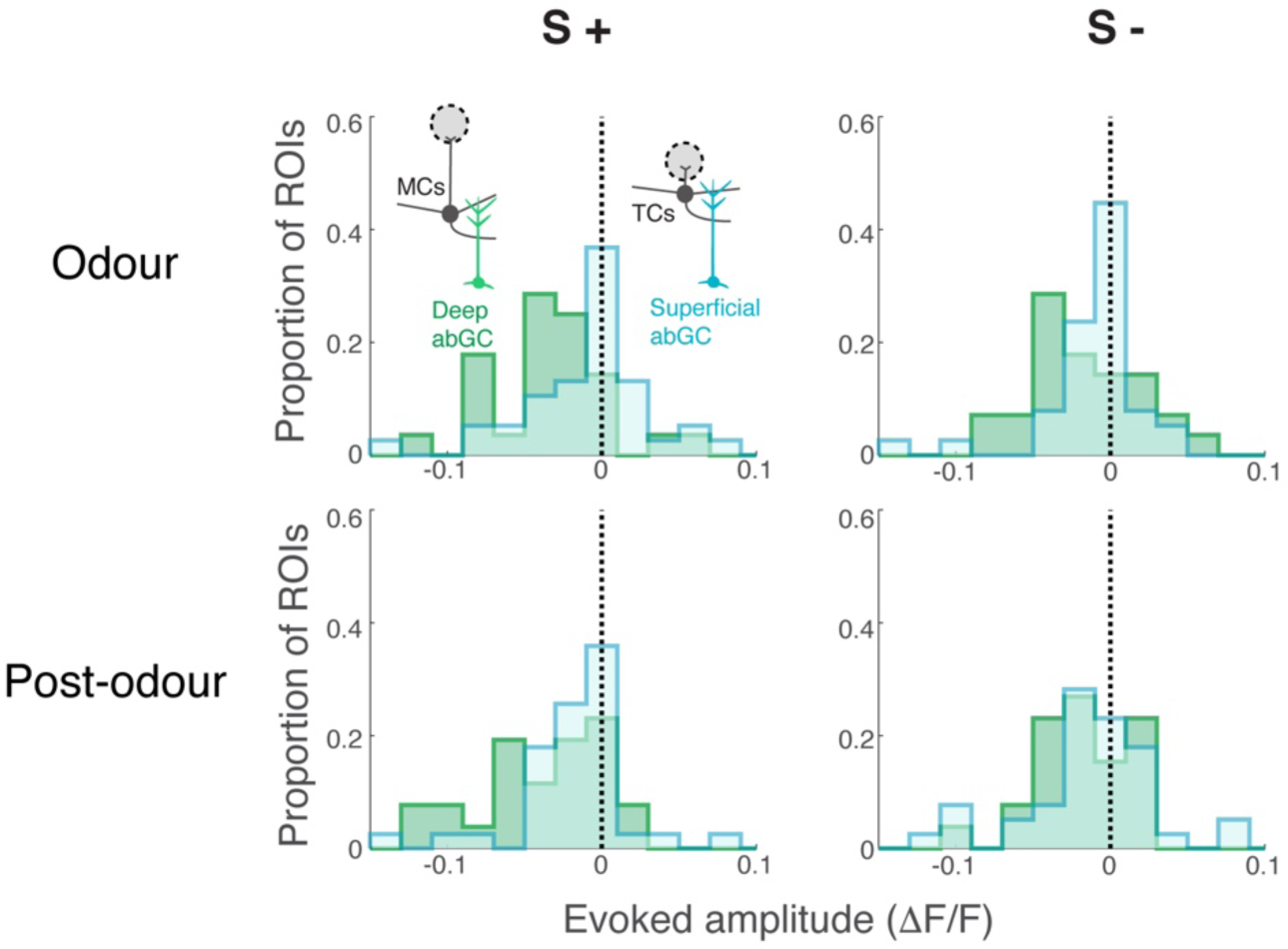
Statistics of evoked responses in depth-resolved adult-born granule cell dendrites. Histograms represent the frequencies of observed fluorescence changes in response to the rewarded odour (S+, left column) and unrewarded odour (S−, right column) for the odour period (top row) and post-odour period (bottom row). Green histograms correspond to data from deep gemmules, and light blue histograms correspond to superficial gemmules.

